# Abundance of Northern Hemisphere tree species declines in the warm and arid regions of their climatic niches

**DOI:** 10.1101/2023.09.04.556202

**Authors:** Julen Astigarraga, Adriane Esquivel-Muelbert, Paloma Ruiz-Benito, Francisco Rodríguez-Sánchez, Miguel A. Zavala, Albert Vilà-Cabrera, Mart-Jan Schelhaas, Georges Kunstler, Christopher W. Woodall, Emil Cienciala, Jonas Dahlgren, Leen Govaere, Louis A. König, Aleksi Lehtonen, Andrzej Talarczyk, Thomas A. M. Pugh

## Abstract

Climate change is expected to drive species towards colder and wetter regions of their distribution with alternative processes such as forest management having the potential to alter species displacements. Here, using data from more than two million monitored trees from 73 widely-distributed species, we quantify changes in tree species abundance across Northern Hemisphere forests and find a widespread decline in abundance across the whole of species’ climatic niches. Yet, our analysis revealed that this decline is heavily influenced by alterations at the stand-level and consequent stand development. Remarkably, when accounting for stand development, our findings show a consistent trend of species abundance optimum shifting towards cold and wet regions within their climatic niches. We provide species-specific information on the direction and magnitude of climate-driven changes in abundance that should be taken into account when designing conservation, management and restoration plans in an era of unprecedented human-caused environmental change.

## Main text

Forests worldwide are largely shaped by human activity (1). Humans have strongly modified forest structure, composition and distribution for millennia through land-use changes and alteration of disturbance regimes (2, 3). In the Northern Hemisphere, the concurrent changes in land-use and intensive forest harvesting have decreased stand age (4) and increased forest area and biomass (5, 6). These human-driven alterations in stand development directly determine tree demographic responses to climate by modifying functional traits related to tree age and size, density-dependent processes, and species interactions (7–10). At the same time, temperature and water availability are major constraints on tree demography (11, 12), and since temperature and water deficit are increasing due to climate change, shifts in species abundance towards relatively colder and wetter regions are expected (13). However, studies tracking shifts in tree species abundance have shown multiple directions in response to climate (14–18), largely because climate responses interact with changes in stand development (19). Thus, a substantial challenge lies in accurately predicting climate-driven shifts in species abundance while accounting for the effects of stand development.

Here, we use harmonized data from national forest inventories from Europe and North America to analyze changes in species abundance across the period 1985-2019 (Mean ± SD census interval = 7 ± 3). For our analyses, we considered more than two million measured trees from 73 widely-distributed species across 126,422 forest inventory plots (Table S1.1). We quantified changes in species abundance (i.e. changes in the number of stems per hectare) in relation to climate and stand development. Specifically, we tested whether and in which direction species abundance is changing across different regions of the species’ climatic niches and whether these changes are modulated by local stand development. We also examined the relationship between species’ climatic niches, measured as the tolerance to cold temperatures and aridity, and the magnitude of climate-driven abundance changes.

We quantified changes in species abundance by partitioning the climatic niche occupied by each species based on minimum temperature and aridity. We found that species abundance changed according to the position of the population in the species’ climatic niche (see Fig. 1A, B as an example for two species with contrasting climatic niches). Despite remarkable variation across species, the number of stems per hectare decreased on average across all climatic regions (Fig 1C). Approximately two-thirds of the species experienced a decline in abundance within each of the climatic regions (Table S3.1, Fig. S3.1 and Fig. S3.2).

**Figure 1.**
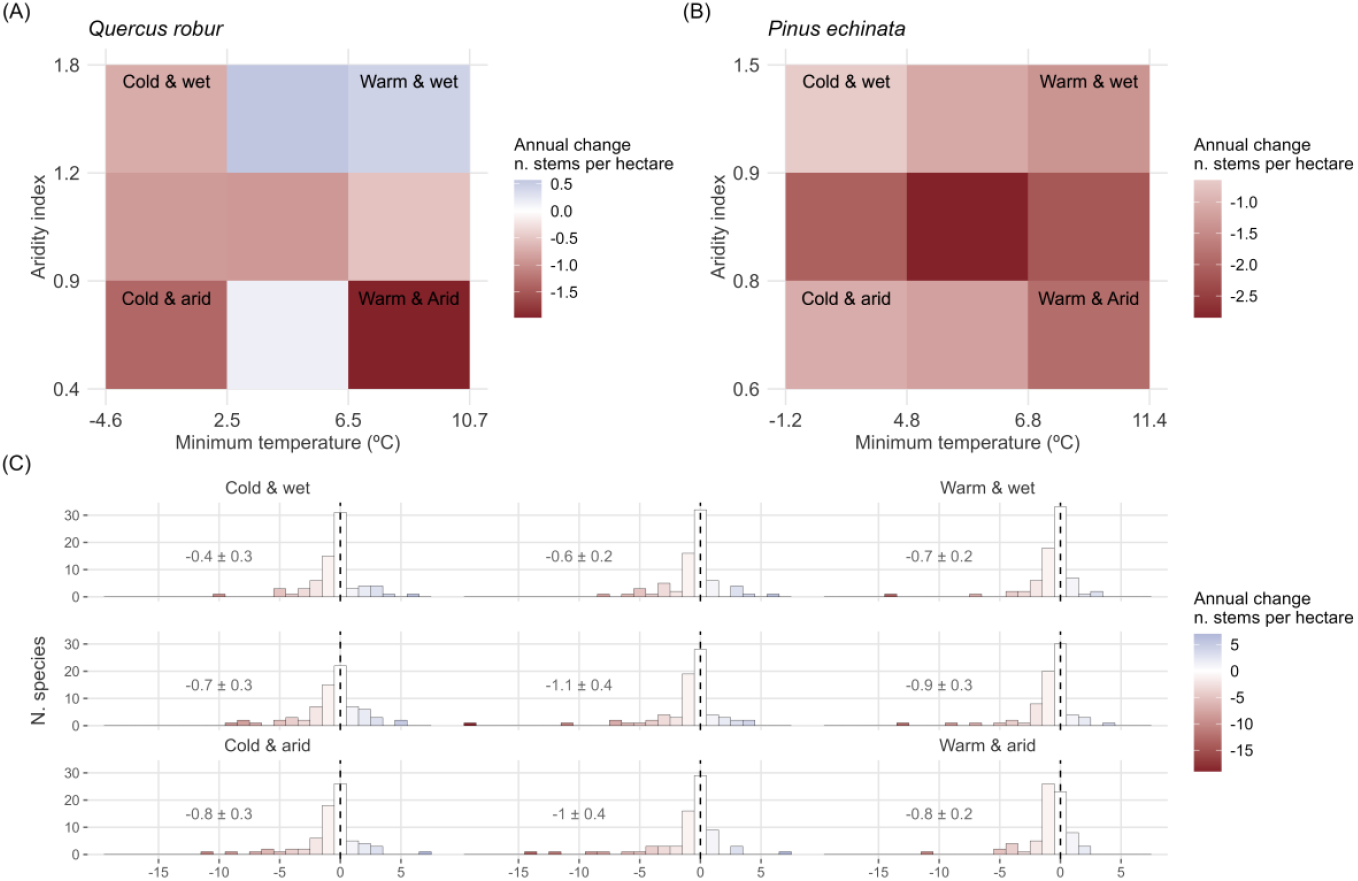
Widespread decline in species abundance across their climatic niches. We partitioned the climatic niche occupied by each species into nine climatic regions defined by the terciles in minimum temperature and aridity across the species’ observed range. For each of the nine regions we calculated the mean change in the annual number of stems per hectare, considering all inventory sites sharing those climatic conditions. Panels in the top row show results for two illustrative species with contrasting climatic niches (A) *Quercus robur* and (B) *Pinus echinata*. (C) Histograms showing the mean change in the annual number of stems per hectare for all 73 species and for each of the nine climatic regions defined by temperature and aridity terciles. Numbers show the overall mean change ± standard error across species, and dashed lines indicate no change. Negative values indicate a decrease in abundance. Note that legends in (A) and (B) are on different scales, and higher values of the aridity index imply less aridity.

The widespread decline in species abundance could result from self-thinning during stand development (20). Alternatively, it could be attributed to varying land-use histories and differing forest management intensities within species’ climatic niches (21, 22). For example, we observed that species mainly distributed in southern Europe have increased in abundance, which is a direct consequence of the abandonment of agricultural activities and traditional forest-use (Fig. S3.1; (23)). Additionally, in central and northern Europe, the decline in abundance of broadleaved species can be attributed to self-thinning and management practices, where broadleaved species have largely been replaced by conifers over the study period. However, it might also indicate that a decrease in species abundance in warm and arid regions is not accompanied by an increase in cold and wet regions, and therefore species are not following climate change in European and U.S. forests (18, 24). Thus, if we aim to quantify the effect of climate change on shifts in species abundance, it is crucial to adjust climate-driven predictions for stand-level changes.

The history of land-use change and forest management may reinforce or complicate shifts in climate-driven species abundance by modifying stand development (21, 22). To account for the interaction between climate and stand development we fitted generalized additive models for each species using a tensor product smoother including this interaction. Using predictive comparisons (25), we quantified the average predictive difference (in number of stems per hectare) in cold and wet (i.e. first quartile of minimum temperature and third quartile of aridity index), median (i.e. second quartiles of minimum temperature and aridity index), and warm and arid regions (i.e. third quartile of minimum temperature and first quartile of aridity index; Fig. 1). In addition, we assessed the effect of stand development by quantifying the average predictive difference in actual, early and late stand development stages (i.e. actual, first and third quartile of stand development, see *Methods*). When adjusting for stand development, we found that species abundance showed larger increases, or smaller decreases, in cold and wet regions compared to warm and arid regions within species’ climatic niches (Fig. 2). For example, *Quercus robur* followed this pattern by increasing in abundance in cold and wet regions of its climatic niche, and decreasing in warm and arid regions (Fig. 2A). Interestingly, in late-development stands, the difference in abundance change between cold and wet and warm and arid regions was more pronounced compared to early-development stands (Fig. 2B). This finding highlights the key role of stand development in shaping species responses to climate (26), and shows that the results depicted in Fig. 1 are largely driven by stand development. Importantly, it shows that the abundance shift towards cold and wet regions is maximized at late-development stands, whereas at earlier stages of stand development the effect of climate shows greater variability between species. This variability in young stands may be a result of the effect of recent disturbances masking any climate effect (27) or the high plasticity of young stands to climate variations (28).This result underlines the importance of focusing on old-growth forests to enhance their persistence in the face of climate change.

**Figure 2.**
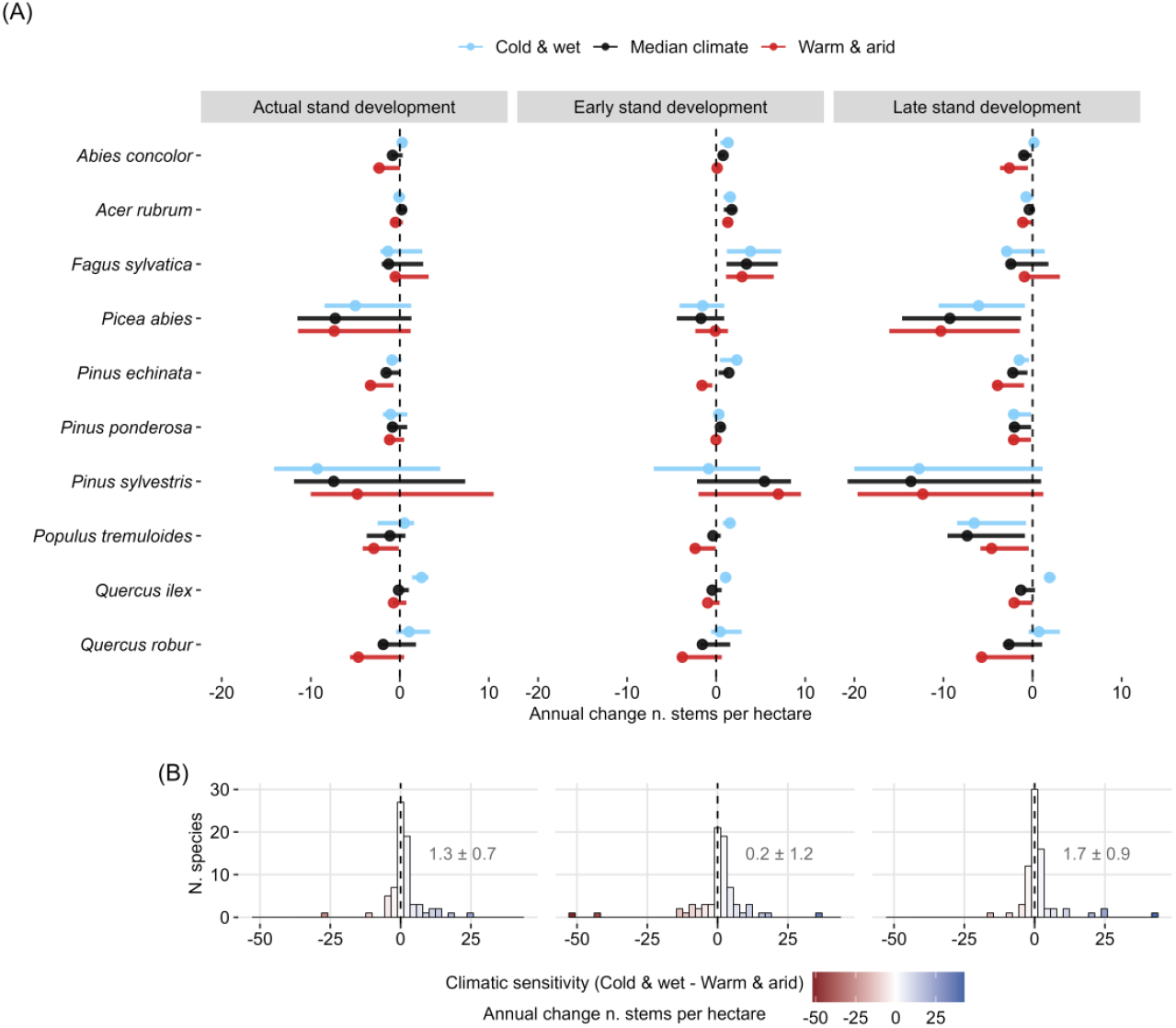
Climate-driven changes in species abundance adjusting for stand development. (A) Predicted changes in species abundance (i.e. annual change in the number of stems per hectare) by species-level models when setting minimum temperature and aridity index in cold and wet (blue), median climate (black), and warm and arid (red) conditions within each species’ climatic niche, and setting stand development in actual, early and late stand development values (see *Methods*). Points indicate mean changes in species abundance and intervals 50% uncertainty, with positive values indicating increases and negative values decreases in abundance over time. The selection of species for representation was made by covering climatic gradients of Europe and the U.S. A full set of results for all species is found in Fig. S3.3, Fig. S3.4 and Table S3.2. (B) Histograms showing the mean climatic sensitivity of each species in actual, early and late stand development conditions. Species’ climatic sensitivity was calculated as the mean difference in the number of stems per hectare between cold and wet regions and warm and arid regions of each species’ climatic niche. Positive values indicate that in cold and wet regions species are gaining more individuals, or losing fewer individuals, than in warm and arid regions. Negative values indicate that cold and wet regions are gaining fewer, or losing more, individuals than warm and arid regions. Numbers on each histogram panel show the overall mean climatic sensitivity ± standard error across species, and dashed lines indicate no difference between cold and wet and warm and arid regions within species.

When determining changes in abundance within species, we found that the majority of species (66% and 60% of species in early- and late-development stands, respectively) showed a significant trend of increasing relative abundance towards cold and wet regions (Table S3.2). Thus, our findings reveal a consistent trend of species abundance shifting towards cold and wet regions, both between and within species across two continents. This could be a consequence of a faster contraction of species ranges in warm and arid regions compared to their expansion in cold and wet regions, potentially exacerbated by more intense forest management in the former (29). Overall, it indicates that the abundance of Northern Hemisphere tree species is declining in the warm and arid regions of their climatic niches, implying heightened vulnerability of populations in these regions to climate change and generalizing previous results at smaller spatial extents (30–32).

The diverse evolutionary histories of species have resulted in a wide range of responses to climate (33). However, identifying key species-level attributes that drive species responses to climate is crucial for predicting their vulnerability to climate change. Thus, we tested whether species’ climatic sensitivity (i.e. mean difference in abundance changes between cold and wet and warm and arid regions of each species’ climatic niche, as shown in Fig. 2B) was related to species’ temperature or aridity niche position using linear mixed effects models. We only fitted the model setting stand development values to late-development stands where the climatic sensitivity of species was found to be larger. We did not find a relation between species’ niche position and their climatic sensitivity (Fig. 3). Hence, species associated with cold and wet climates did not show larger climatic sensitivity than species inhabiting warm and arid climates. Instead, the positive average species’ climatic sensitivity reported in Fig. 2 was maintained over the whole range of climatic niche spaces. On average, species showed larger increases in abundance, or lower decreases, in relatively cold and wet climatic conditions, regardless of their temperature or aridity affinities. This suggests that temperature or aridity tolerance does not render a species safe from the effects of climate change (34).

**Figure 3.**
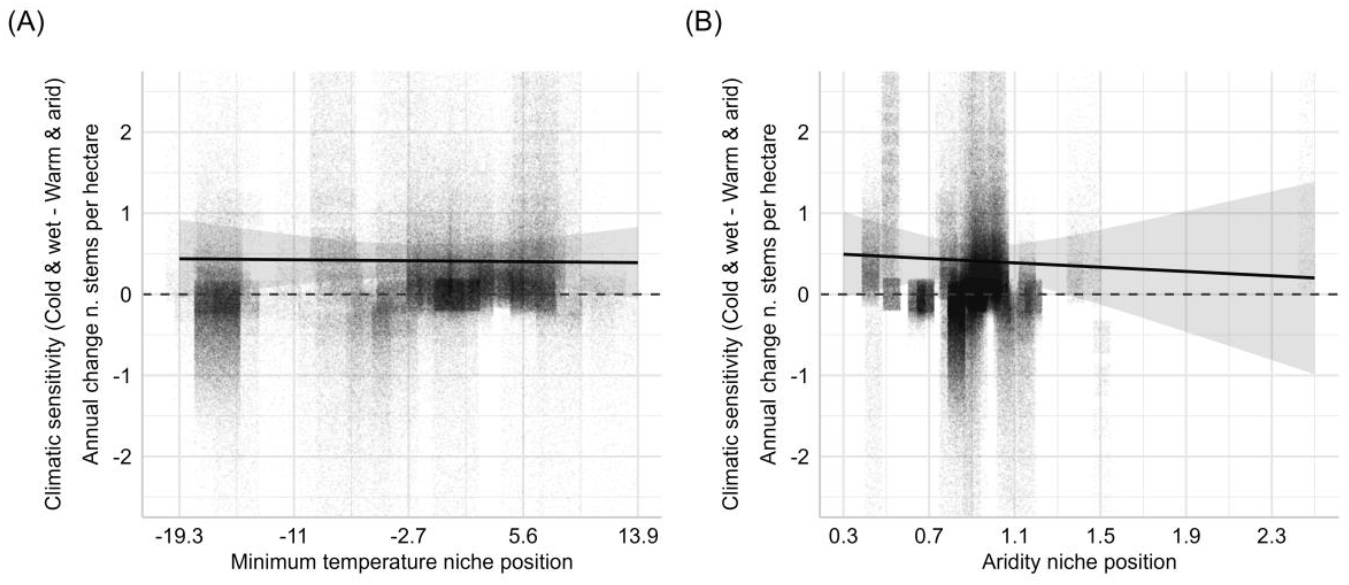
Shifts in species abundance towards cold and wet regions in relation to species’ niche position in late-development stand. Predicted species’ climatic sensitivity along (A) minimum temperature niche position and (B) aridity niche position. Positive values indicate that in cold and wet regions species are gaining more individuals, or losing fewer individuals, than in warm and arid regions; negative values indicate the opposite. The shaded areas show the confidence intervals for the regression slope based on ± 1 standard error, and points are the climatic sensitivity of each species on each plot where the species was present. To improve pattern visibility, Y-axis limits are zoomed in to -2.5 and 2.5 and species’ climatic sensitivity points are jittered.

Overall, we observed a displacement in abundance optimum of Northern Hemisphere tree species towards cold and wet regions of their climatic niches that could jeopardize ecosystem functioning and human well-being (35). The shift in abundance optimum is happening both between and within species but it was mainly observed in late-development stands. These findings highlight the extent to which disturbances that alter stand development are pervasive in shaping climate-driven changes in species abundance (36). The generally positive climatic sensitivity displayed by species with different climatic niches indicates that no forest region is clearly more resilient against the niche-shifting effects of climate change. However, the significant number of species that show negative climatic sensitivities (34% and 40% in early- and late-development stands, respectively, Table S3.2) suggests that factors other than climate may be influencing the abundance of these species in cold and wet regions or that these particular species are relatively more resilient to ongoing climate change. Plausibly these species occupy a subset of the range of plant functional strategies that can thrive within a specific climatic niche, which would imply that whilst climate niche is not an indicator of climatic sensitivity, the functional strategy of species is likely to play a significant role (37, 38). It remains unclear whether these species would continue to show benefits on their warm and arid region once migration of species typically associated with warmer and more arid regions is completed and new species assemblages emerge. Therefore, one of the next critical frontiers is to account for novel competitive interactions to accurately predict species responses to climate change (39). By determining climate-driven species abundance changes, our study represents a crucial step towards reducing the uncertainty surrounding the future climate risk faced by forests (40). In addition, the quantification of species’ climatic sensitivity holds substantial implications for conservation planning, as well as for identifying which species can best withstand the escalating impacts of climate change to support forest restoration and adaptation.

## Materials and Methods

### Forest inventory data

We calculated changes in species abundance using harmonized forest inventory data from seven European countries and the continental United States, with censuses spanning between 1985-2019 (Mean ± SD census interval = 7 ± 3; Table S1). For all countries, we analyzed the last two available consecutive forest inventory censuses with a census interval ≥ 4 years. For all tree species, we excluded plots where the species was missing (i.e. zero stems were recorded) in the first census, as our focus was on assessing changes in abundance in regions where species were already present. We also excluded plots where all trees were missing in the subsequent census to mitigate potential management-related impacts like clear-cutting. In the end, we used a total of 126,422 plots. We calculated changes in species abundance as changes in the annual number of stems per hectare between censuses. In plots where not all trees were measured in all subplots (e.g. variable radius plots) we generated stochastic stem counts from a one-truncated poisson distribution with λ equal to the number of stems extrapolated to the plot area using *extraDistr* R package (41). To select the most widely-distributed species, we aggregated the data into 0.1º × 0.1º grid cells (i.e. ∼11.1 km at equator) and we only considered native species to Europe and the U.S. that have a good coverage in our dataset (i.e. species with ≥ 50,000 individuals of > 12.7 cm in diameter at breast height and that were present in ≥ 500 cells and ≥ 1,000 plots in our dataset). In total we selected 73 species (Table S1.1 and S3.2).

### Species’ climatic niches

We quantified the climatic niche of each species using the climatological mean minimum temperature and the aridity index. To characterize minimum temperature, we used the mean temperature of the coldest quarter obtained from WorldClim 2 (42). Aridity was obtained from Global Aridity Index and Potential Evapo-Transpiration (ET0) Climate Database v2 (43). This variable is based on the Global Aridity Index and it is calculated by dividing the mean annual precipitation by the mean annual reference evapo-transpiration, with its values increasing for wetter conditions. Both datasets provide 30 arc-seconds spatial resolution (∼ 1km at equator) for the 1970-2000 period.

We characterized species minimum temperature and aridity niche position using species chorological maps for European species (44) and Little’s range maps for U.S. species (45, 46). We quantified species minimum temperature niche position and aridity niche position with the mean minimum temperature and mean aridity index across the whole species’ range, respectively.

### Stand development

Basal area is commonly used as a proxy of stand development (e.g. 26), as basal area increases during stand development (20). However, the maximum stand basal area (i.e. that expected to be observed in late-development stands) depends on climate and soil conditions (47–49). Thus, we calculated stand development based on the basal area of each plot with respect to the maximum basal area found in plots with similar climate and soil characteristics.

First, we created clusters of plots with similar climate and soil characteristics along Europe and the U.S. considering key factors driving basal area increment: annual mean temperature, annual precipitation (ln scale) and nitrogen availability (47–49). Climatological mean annual temperature and precipitation at 1 km^2^ were obtained from WorldClim 2 (42) and nitrogen availability in 0-30 cm at 250 m from SoilGrids v2 (50). Since nitrogen between 0 and 30 cm was obtained from different layer thicknesses (0-5, 5-15 and 15-30 cm), we calculated the weighted mean of nitrogen adjusting for the thickness.

Second, we computed the k-means clustering using *stats* R package (51) for values of *k* ranging from 10 to 100 clusters with increments of 10. For each *k*, we calculated the total within-cluster sum of squares and using the elbow method we identified the optimum number of clusters (Fig. S2.1 and S2.2; (52)). We selected 40 clusters in both Europe and the U.S. with a compactness (i.e. the similarity of the plots within the same cluster) of 95.5% and 96.2% in Europe and the U.S., respectively. Additionally, to ensure that the selection of 40 clusters was optimal, using the same sequence from 10 to 100 with an increment of 10, we analyzed the median cluster size and the minimum cluster size (Fig. S2.1 and S2.2). We also confirmed that the coefficient of variation of cluster precipitation (ln scale), mean temperature and nitrogen availability followed a similar pattern to that of the within-cluster sum of squares (Fig. S2.1 and S2.2). Furthermore, we observed that the coefficient of variation of basal area remained high regardless of the number of clusters, suggesting a high variability of stand development stages within the cluster. Once the number of clusters was set to 40, we validated it using the silhouette method with an average silhouette width 0.26 and 0.29 in Europe and the U.S., respectively (Fig. S2.3 and S2.4).

Finally, for each of the forest inventory plots we obtained a stand development value ranging from 0 to 1 by dividing the basal area of each plot to the maximum basal area of its corresponding cluster (i.e. 95^th^ percentile of the basal area of the cluster to which this plot belonged). For plots exceeding the 95^th^ percentile of the basal area of the cluster we assigned a value of 1.

### Analyses

First, we determined how changes in species abundance varied across each species’ climatic niche. For this aim, we divided the climatic niche of each species into nine climatic regions. These regions were defined by the terciles in minimum temperature and aridity across the species’ observed range. Specifically, populations were categorized based on their location within the 0-33%, 33-66%, or 66-100% range of each species’ distribution in our dataset. This approach allowed us to quantify a climatic niche that ranged from cold and wet to warm and arid regions.

For each of the nine regions we calculated the mean change in the annual number of stems per hectare across all inventory sites sharing those climatic conditions. Then we calculated the mean change in number of stems per hectare for each of the nine climatic regions, averaging values from all species. Changes in abundance across species’ climatic niches were visualized with *terra* and *tidyverse* R packages (53, 54) and all analyses were performed using R Statistical Software (v4.2.0; (51)).

Second, we quantified the effect of climate and stand development on species abundance across each species’ climatic niche by fitting generalized additive models (GAM; (55); (56)) to each species separately using *mgcv* R package (57). We modeled the number of stems in the second census following a negative binomial distribution with a tensor product smoother of aridity, minimum temperature and stand development of each plot. The basis for the tensor product smoothers were cubic regression splines. We also included the interaction of the natural logarithm of the number of stems in the first census and the census interval as fixed effects to adjust for the initial number of stems in the plot and the number of years elapsed between censuses, respectively. In addition, we included the country in which each plot was measured to adjust for different sampling methods among countries, and an offset of the natural logarithm of plot area to adjust for different plot areas. We diagnosed each species’ GAM fit, checking the residuals plots using *mgcv, gratia* and *DHARMa* R packages (57–59). We also checked the relationship between the response and main effects with *visreg* R package (60), estimated smoothers using *gratia* R package (58), the performance of the model using *performance* R package (61) and evaluating predictions on the observed data.

We evaluated the effects of climate and stand development on species abundance using average predictive comparisons (25). To calculate the effect of climate, we predicted changes in abundance in different conditions: first setting minimum temperature in the first quartile and aridity index in the third quartile (i.e. cold and wet), second by setting minimum temperature and aridity index in the second quartiles (i.e. median climate), and finally setting minimum temperature in the third quartile and aridity index in the first quartile (i.e. warm and arid) of each species’ climatic niche. Each of these three predictions was repeated three times: (i) using the actual stand development values in the plot; (ii) setting stand development values to the first quartile when considering all species together (i.e. early stand development); and (iii) setting stand development values to the third quartile when considering all species together (i.e. late stand development). Then, we obtained annual changes in species abundance per hectare as the difference between the predictions of each model and the number of stems in the first census, extrapolated to the hectare and dividing by the years elapsed between censuses. The climatic sensitivity of each species was calculated as the difference in the annual change in the number of stems per hectare between cold and wet regions, and warm and arid regions in actual, early and late stand development. Positive values indicated that in cold and wet regions species were gaining more individuals, or losing fewer individuals, than in warm and arid regions; negative values indicated the opposite. We visualized changes in species abundance along climatic and stand development gradients using *ggdist* and *tidyverse* R packages (54, 62).

Finally, we calculated the effect of species’ minimum temperature and aridity niche position on species’ climatic sensitivity. We also used the difference in the annual change in the number of stems per hectare between cold and wet regions and warm and arid regions to quantify species’ climatic sensitivity. We fitted a linear mixed effects model assuming a normal distribution using *lme4* package (63). We included species minimum temperature niche position and aridity niche position as fixed effects, and species identity and plot identity as random effects. We fitted the model setting stand development values to the third quartile when considering all species together (i.e. late-development stands), where climatic sensitivity was found to be larger. Both fixed predictors were standardized before being included in the models. We diagnosed each model fit, checking the residuals plots using *DHARMa* R package (59) and predictions were visualized using *ggeffects* and *tidyverse* R packages (54, 64).

## Acknowledgments

JA, PRB, MAZ, AVC and TAMP acknowledge funding from the CLIMB-FOREST Horizon Europe Project (No 101059888) that was funded by the European Union. JA, PRB and MAZ were funded by the Spanish Ministry of Science and Innovation (subproject LARGE, No PID2021-123675OB-C41). TAMP, AEM, MJS and JA acknowledge funding from the European Research Council (ERC) under the European Union’s Horizon 2020 research and innovation programme (grant agreement No 758873, TreeMort). This is a contribution to the strategic research areas BECC and MERGE and to the Nature-based Future Solutions profile area at Lund University. JA was supported by a FPI fellowship of the Department of Education of the Basque Government, AEM by the UKRI TreeScapes MEMBRA, FRS by VI Plan Propio de Investigación of Universidad de Sevilla (VI PPIT-US), FEDER 2014-2020 and Consejería de Economía, Conocimiento, Empresas y Universidad of Junta de Andalucía (grant US-1381388, Universidad de Sevilla), and AVC by a Juan de la Cierva-Incorporación fellowship (IJC2018-038508-I) from the Ministry of Science and Innovation (Spain). We extend our gratitude to the institutions and teams behind the Flemish Forest Inventory, CzechTerra Landscape Inventory, Finnish Forest Health Monitoring Network, Dutch Forest Inventory, Polish National Forest Inventory, Spanish National Forest Inventory, Swedish National Forest Inventory and the Forest Inventory and Analysis (FIA) research program. The Spanish National Forest Inventory is available thanks to the Ministry for the Ecological Transition and the Demographic Challenge (MITECO) and the Polish National Forest Inventory was funded by the State Forests Holding (Lasy Państwowe). We thank the team behind the Global Forest Dynamics database, initiated by the TreeMort project, on which this study is based. We also thank V. Cruz-Alonso, J. M. Serra-Diaz and F. Lloret for the interesting discussion on the manuscript and their useful comments to improve it.

