## Supporting information for "Abundance of Northern Hemisphere tree species declines in the warm and arid regions of their climatic niches"

**Appendix S1. National forest inventories**

**Appendix S2. Stand development**

**Appendix S3. Further analyses**

**SI References**

**Appendix S1. National forest inventories**

**Table S1.1.** Plot type, mean plot area in hectares, minimum census year, maximum census year, mean census interval, number of species and number of plots in each of the national forest inventories analyzed.

| Country | Plot type | Mean plot area (ha) | Minimum census year | Maximum census year | Mean census interval | No. species | No. plots |
|---|---|---|---|---|---|---|---|
| Belgium (Flanders) (1) | variable radius & concentric circles | 0.10 | 1994 | 2019 | 12 | 10 | 1404 |
| Czech Republic (CzechTerra) (2) | concentric circles | 0.05 | 2008 | 2015 | 6 | 11 | 545 |
| Finland (forest health monitoring) (3) | variable radius | 0.03 | 1985 | 1995 | 10 | 5 | 1863 |
| Netherlands (4) | concentric circles | 0.04 | 2001 | 2019 | 7 | 11 | 1142 |
| Poland (5, 6) | concentric circles | 0.03 | 2005 | 2014 | 5 | 12 | 20124 |
| Spain (7, 8) | variable radius | 0.20 | 1986 | 2008 | 11 | 16 | 38681 |
| Sweden (9) | concentric circles | 0.03 | 2003 | 2017 | 5 | 9 | 11705 |
| U.S. (10, 11) | circular | 0.07 | 1998 | 2018 | 6 | 55 | 50958 |

### Appendix S2. Stand development

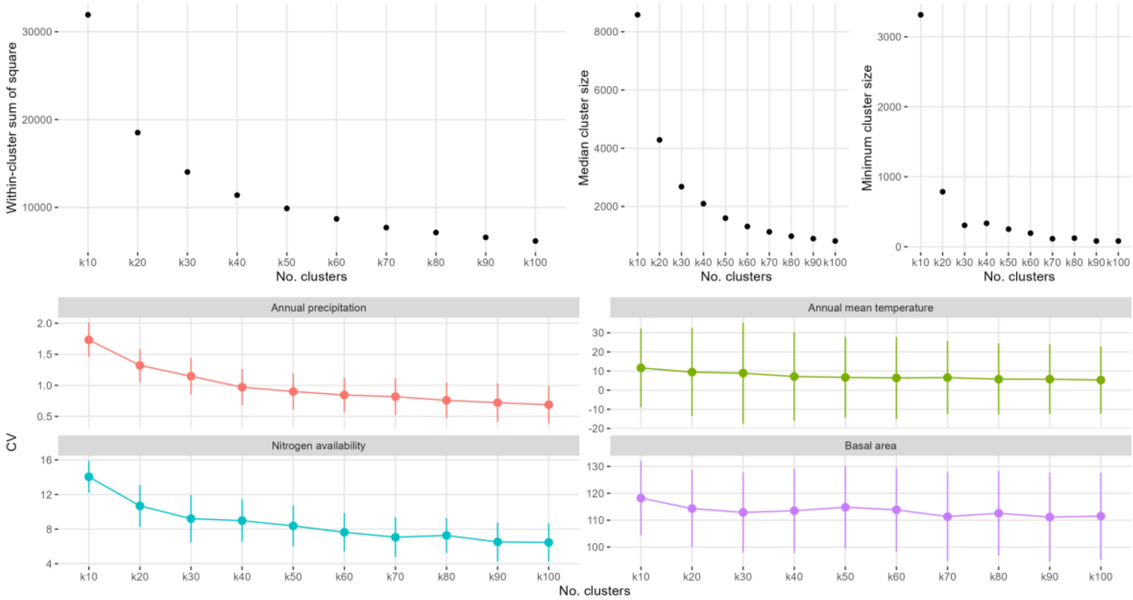


**Figure S2.1.** Metrics for determining the appropriate number of clusters (k = 10, 20, 30, 40, 50, 60, 70, 80, 90 and 100 clusters) for characterizing stand development based on the k-means clustering in Europe. Top panel: within sum of squares, median cluster size and minimum cluster size vs. the number of clusters. Bottom panel: the coefficient of variation of cluster basal area, precipitation (ln scale), mean temperature and nitrogen availability vs. the number of clusters. In the bottom panel, the points and vertical lines indicate the median value and the standard deviation of the coefficient of variation of each cluster, respectively, for each variable analyzed.

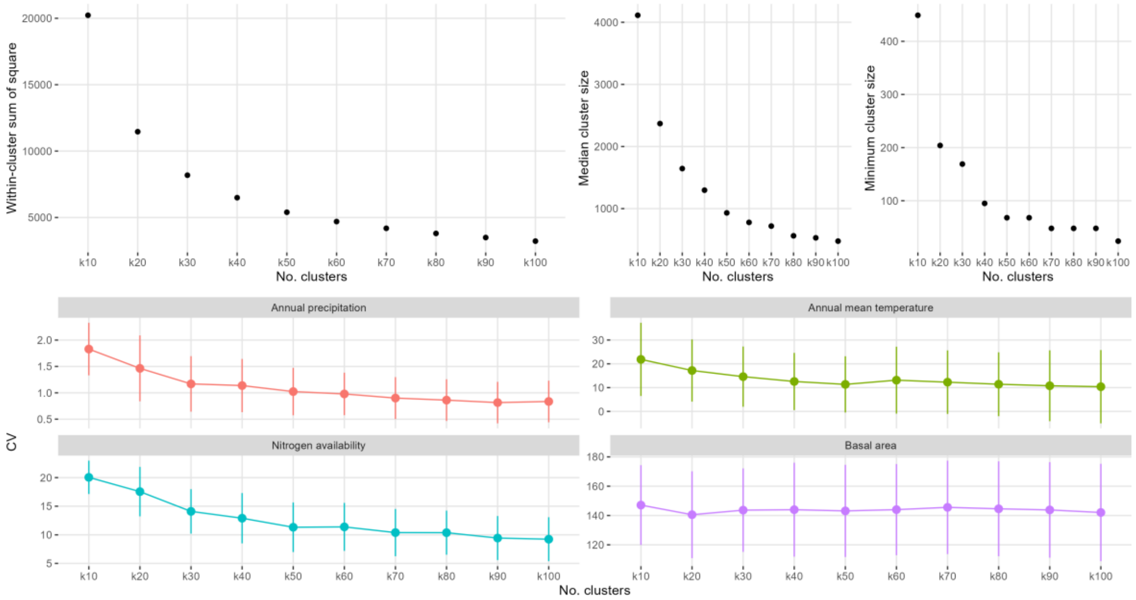


**Figure S2.2.** Metrics for determining the appropriate number of clusters (k = 10, 20, 30, 40, 50, 60, 70, 80, 90 and 100 clusters) for characterizing stand development based on the k-means clustering in the U.S. Top panel: within sum of squares, median cluster size and minimum cluster size vs. the number of clusters. Bottom panel: the coefficient of variation of cluster basal area, precipitation (ln scale), mean temperature and nitrogen availability vs. the number of clusters. In the bottom panel, the points and vertical lines indicate the median value and the standard deviation of the coefficient of variation of each cluster, respectively, for each variable analyzed.

19

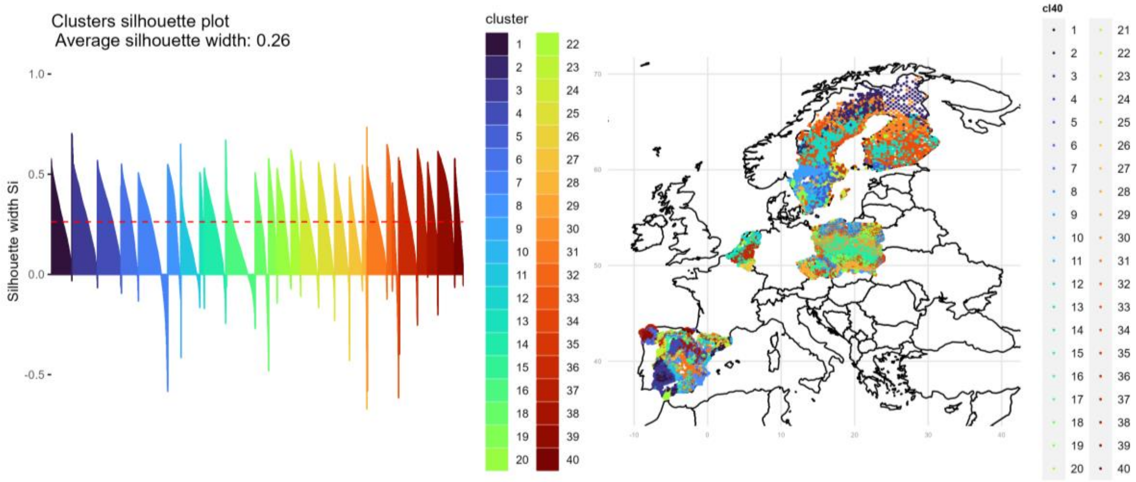


**Figure S2.3.** Cluster validation in Europe based on clusters silhouette plot using 40 clusters. As shown by the silhouette plot and the average silhouette coefficient, most of the silhouette coefficients are positive, indicating that the observations have been placed in the correct group.

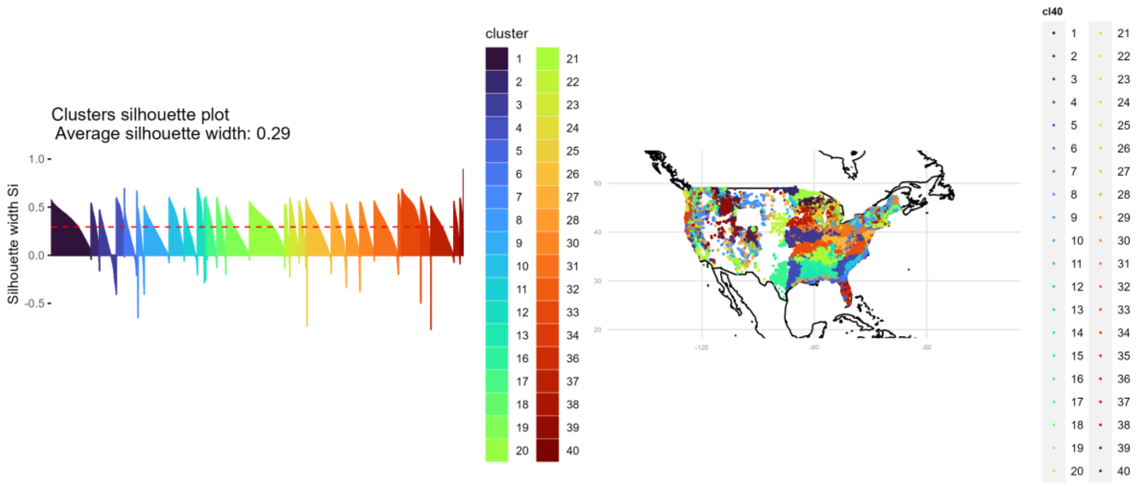


**Figure S2.4.** Cluster validation in the U.S. based on clusters silhouette plot using 40 clusters. As shown by the silhouette plot and the average silhouette coefficient, most of the silhouette coefficients are positive, indicating that the observations have been placed in the correct group.

### Appendix S3. Further analyses

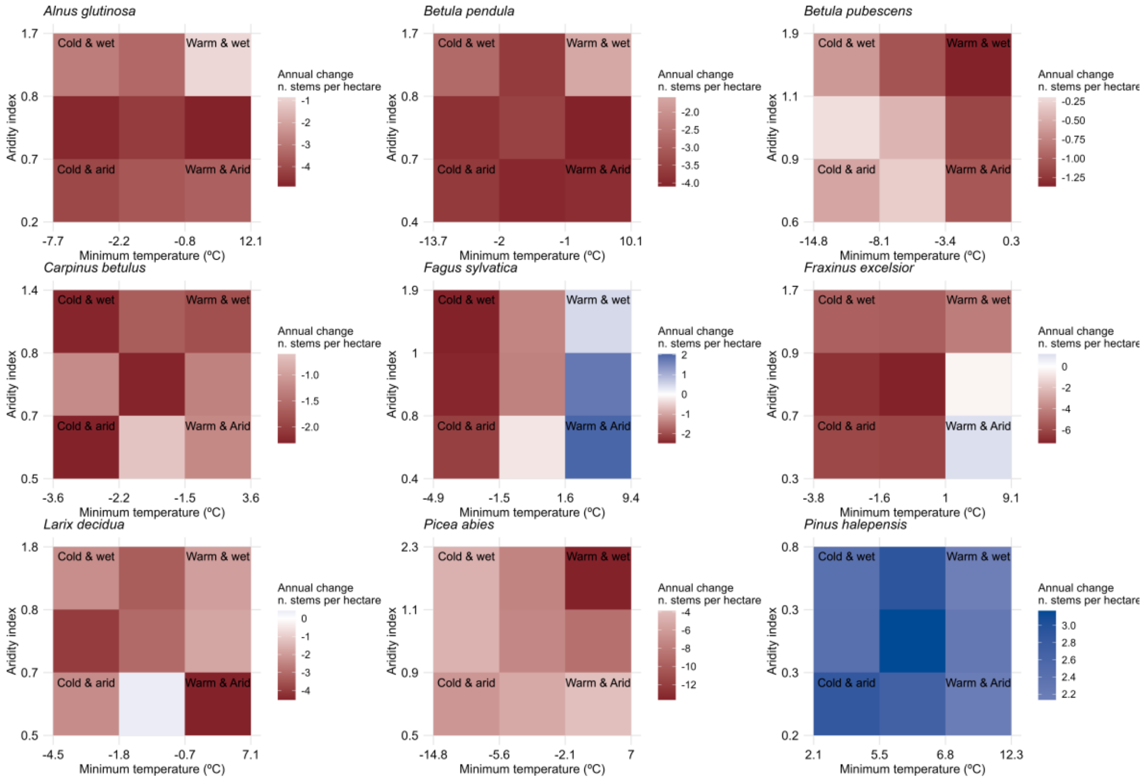


**Figure S3.1.** Mean changes in the annual number of stems per hectare across European species' climatic niche. Note that legends are on different scales.

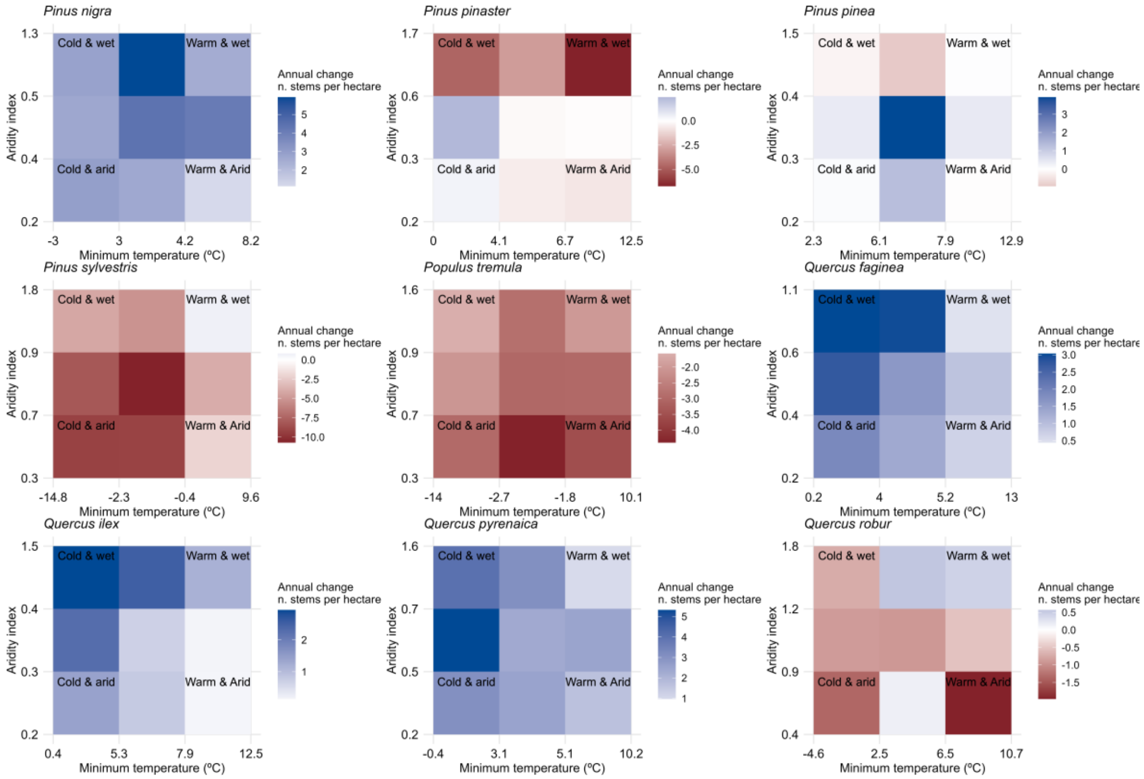


**Figure S3.1 (cotinuation).** Mean changes in the annual number of stems per hectare across European species' climatic niche. Note that legends are on different scales.

25

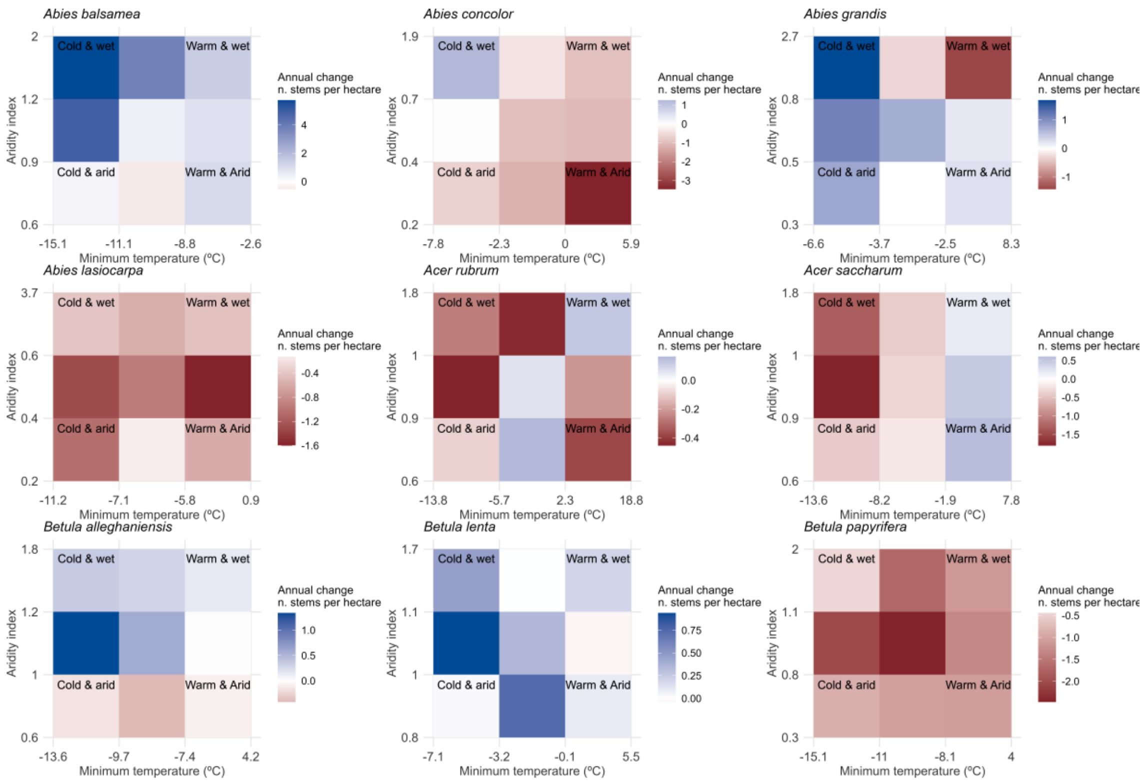


**Figure S3.2.** Mean changes in the annual number of stems per hectare across U.S. species' climatic niche. Note that legends are on different scales.

26

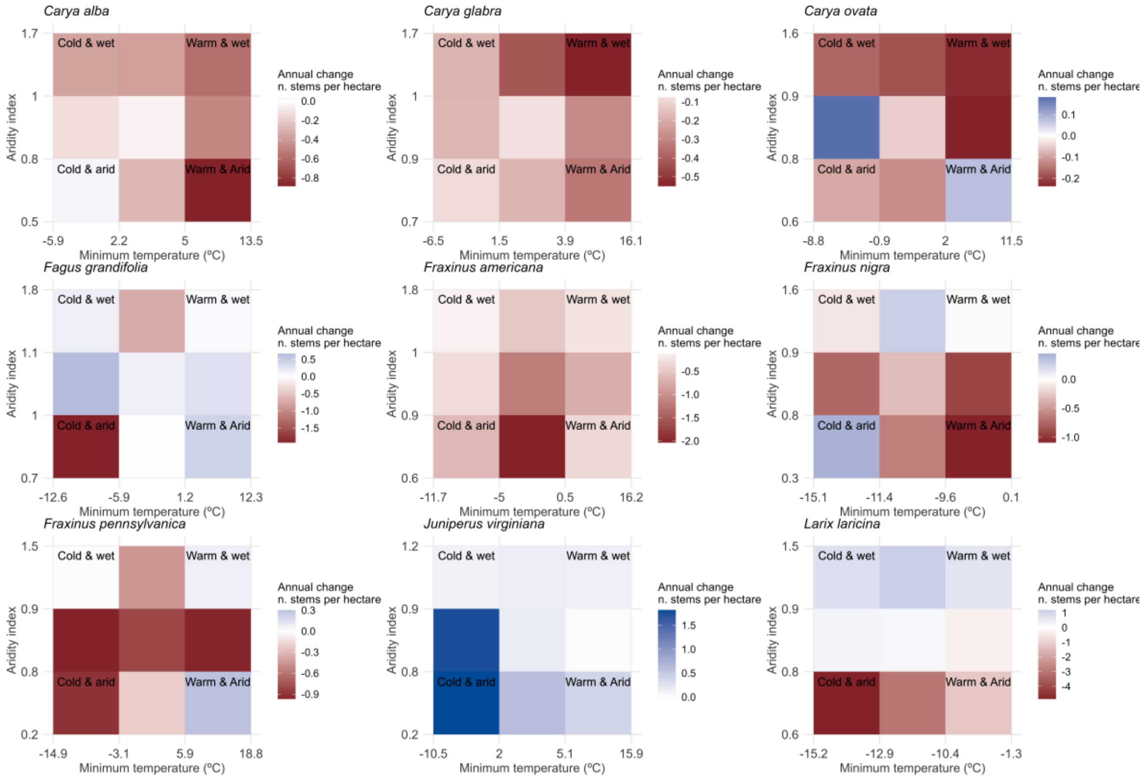


**Figure S3.2 (cotinuation).** Mean changes in the annual number of stems per hectare across U.S. species' climatic niche. Note that legends are on different scales.

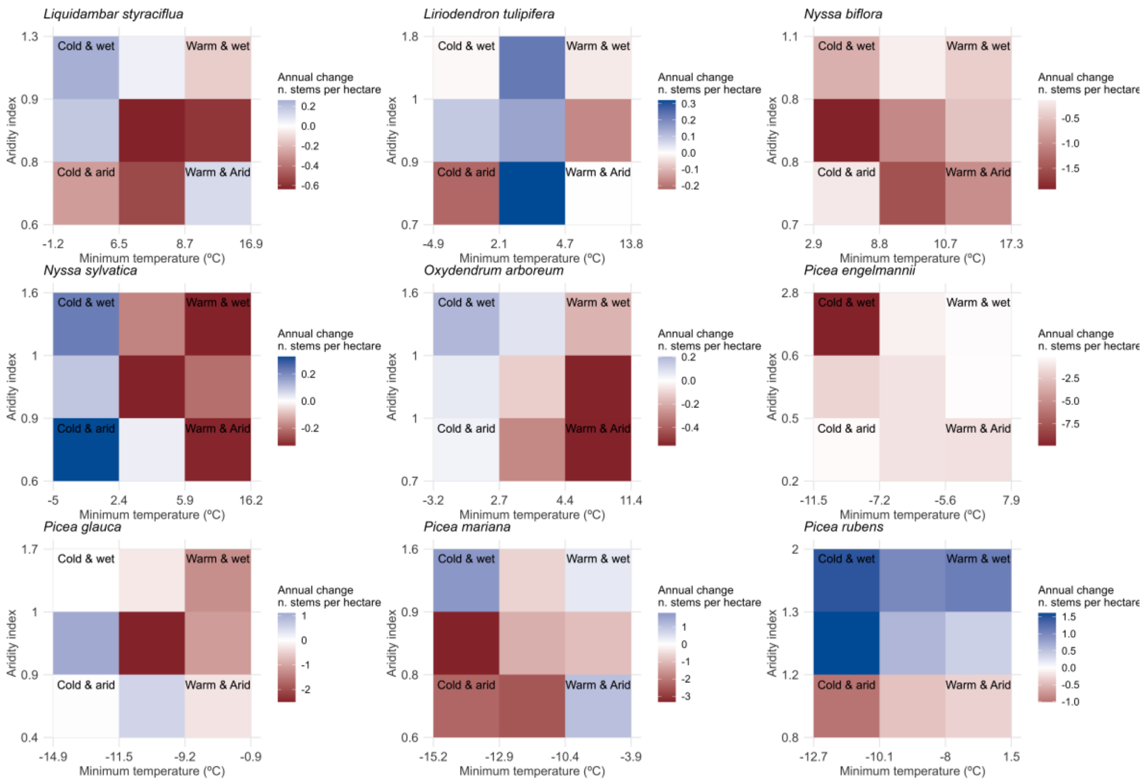


**Figure S3.2 (cotinuation).** Mean changes in the annual number of stems per hectare across U.S. species' climatic niche. Note that legends are on different scales.

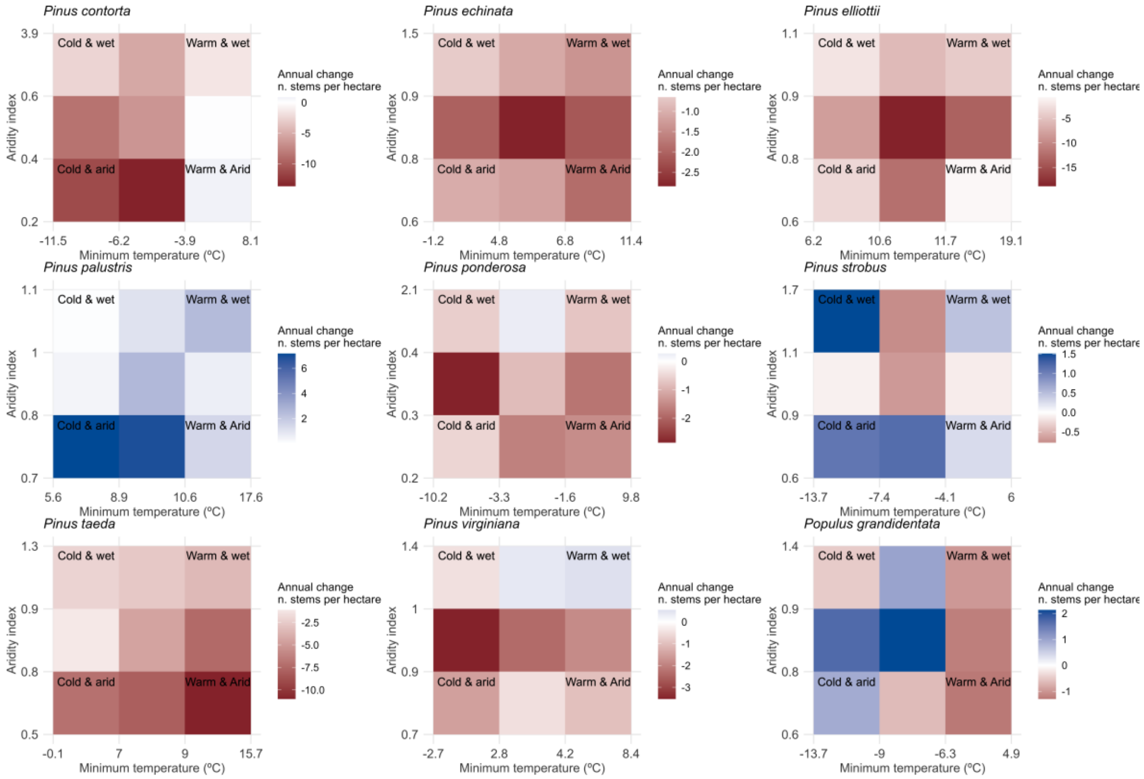


**Figure S3.2 (cotinuation).** Mean changes in the annual number of stems per hectare across U.S. species' climatic niche. Note that legends are on different scales.

29

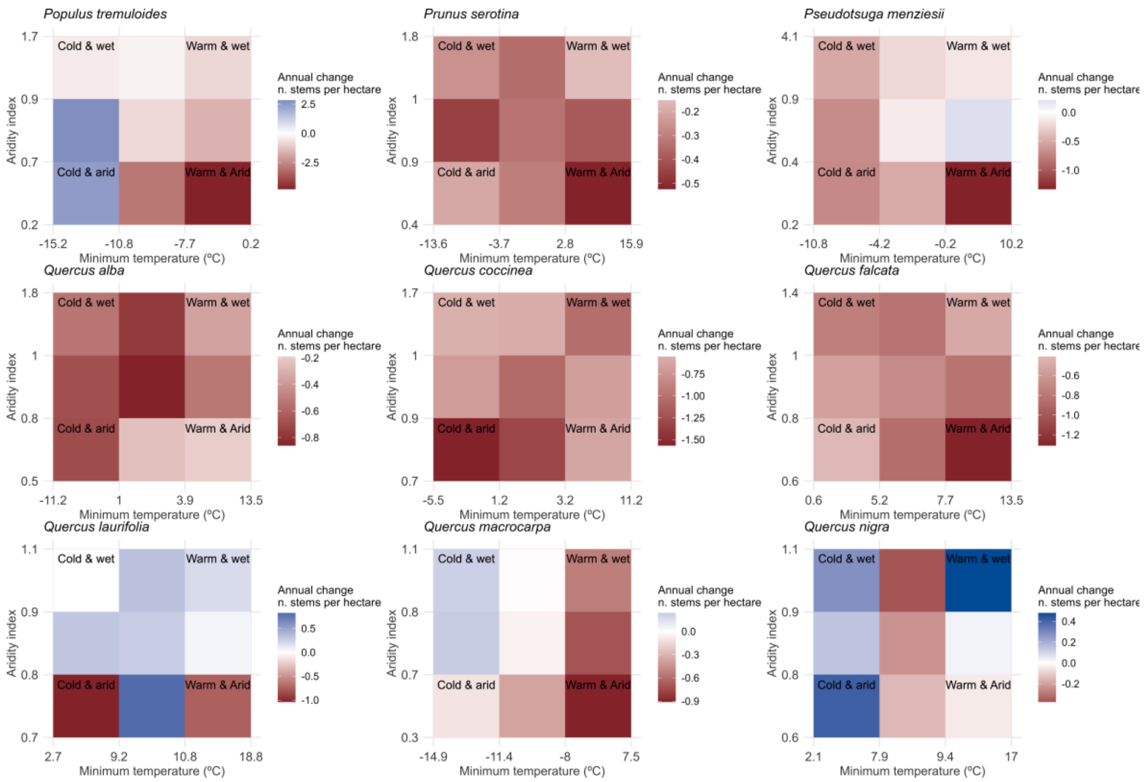


**Figure S3.2 (cotinuation).** Mean changes in the annual number of stems per hectare across U.S. species' climatic niche. Note that legends are on different scales.

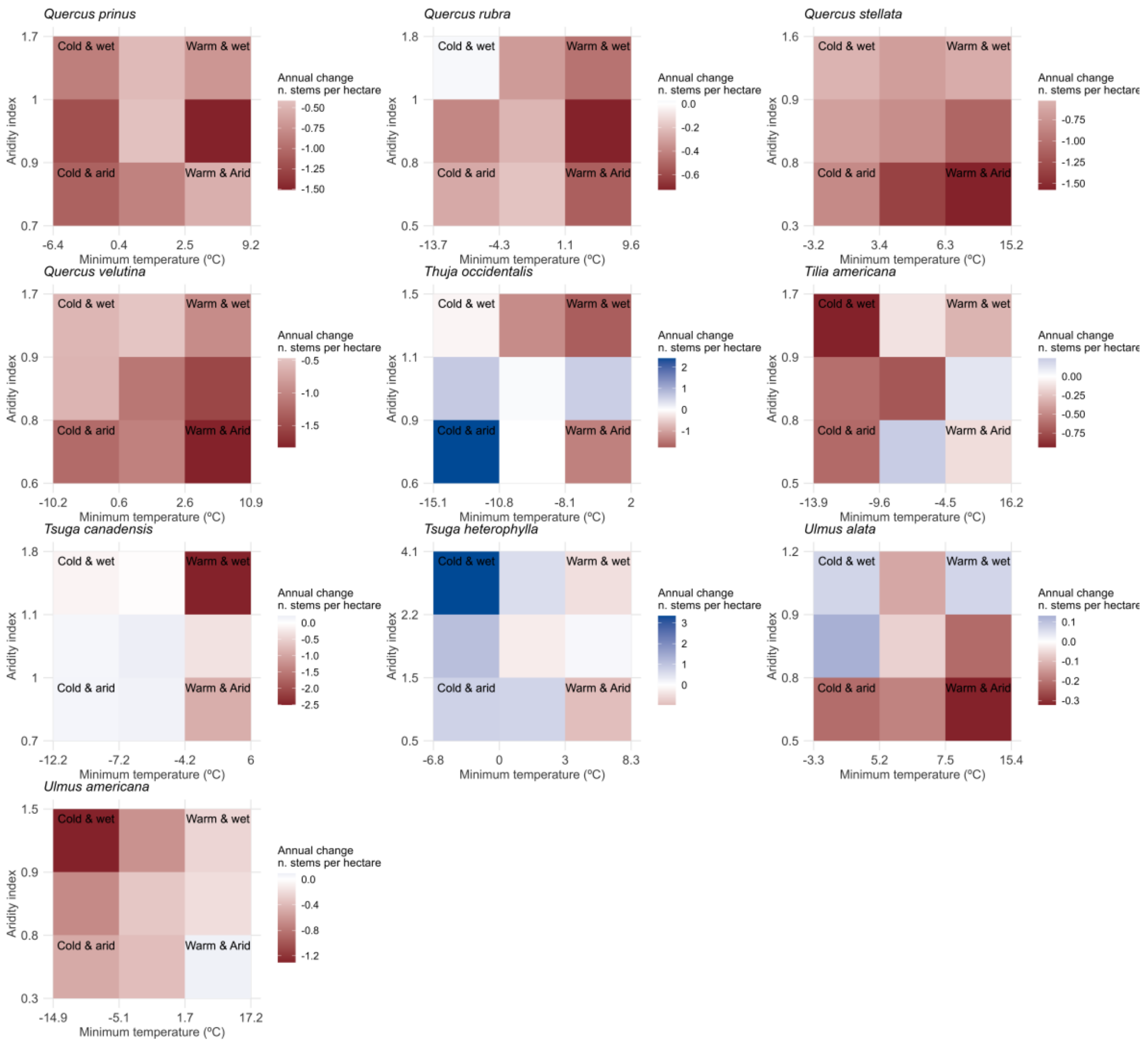


**Figure S3.2 (cotinuation).** Mean changes in the annual number of stems per hectare across U.S. species' climatic niche. Note that legends are on different scales.

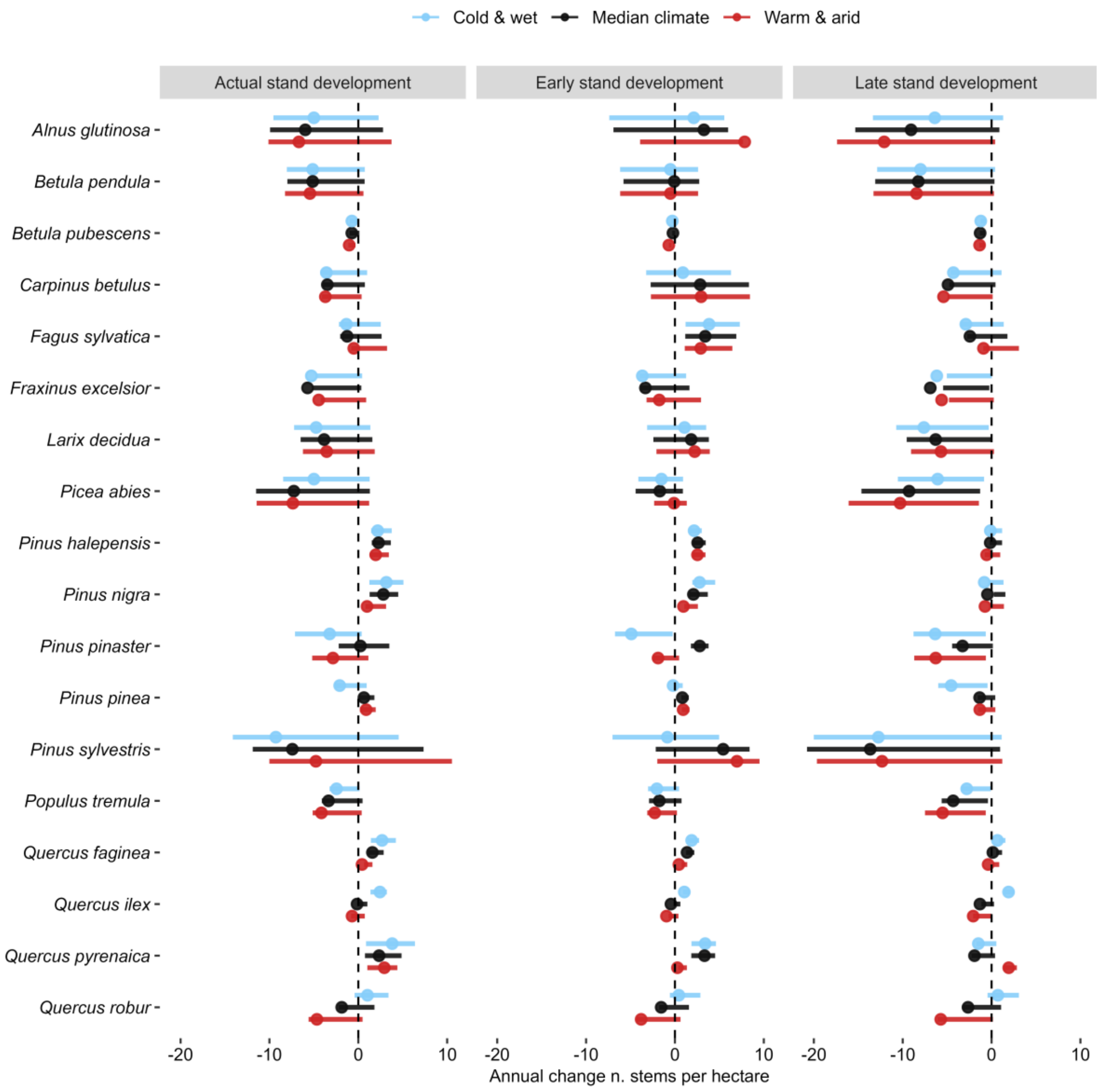


**Figure S3.3.** Predicted changes in European species abundance (i.e. annual changes in the number of stems per hectare) by species-level models when setting minimum temperature and aridity index in cold and wet (blue), median climate (black), and warm and arid (red) conditions within each species' climatic niche, and setting stand development in actual, early and late stand development (see *Methods*). Points indicate mean changes in species abundance and intervals 50% uncertainty.

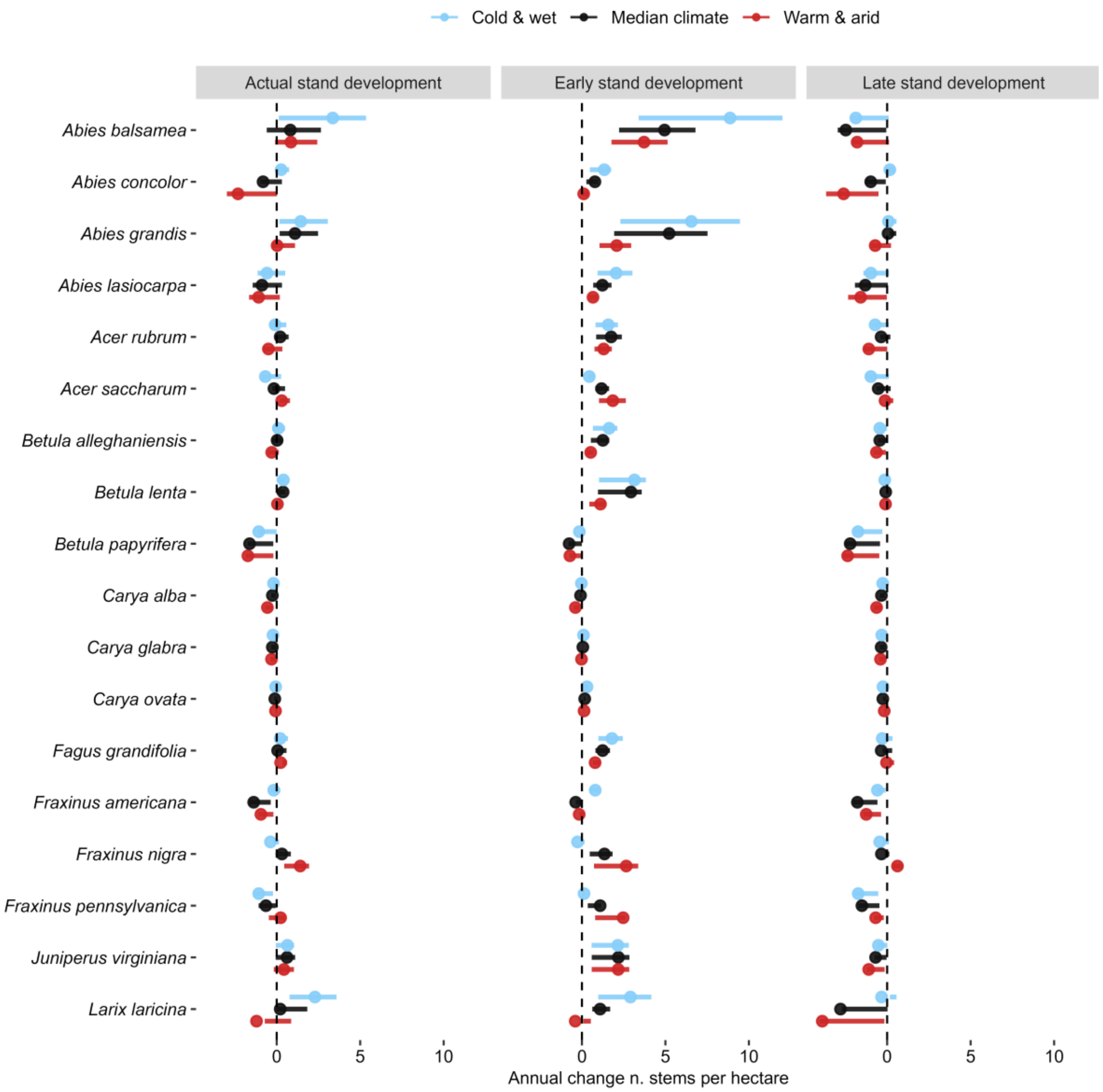


**Figure S3.4.** Predicted changes in U.S. species abundance (i.e. annual changes in the number of stems per hectare) by species-level models when setting minimum temperature and aridity index in cold and wet (blue), median climate (black), and warm and arid (red) conditions within each species' climatic niche, and setting stand development in actual, early and late stand development (see *Methods*). Points indicate mean changes in species abundance and intervals 50% uncertainty.

33

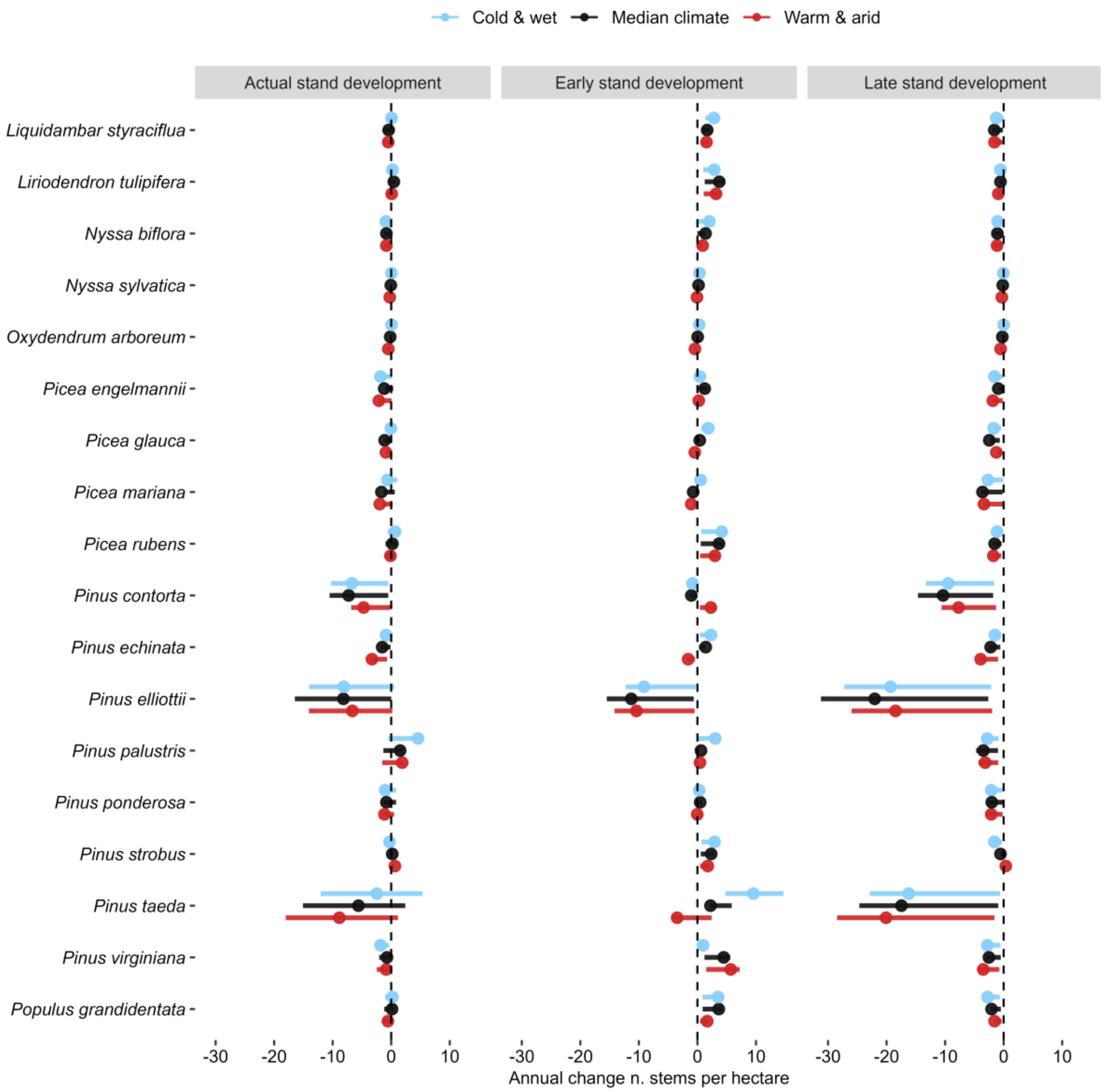


**Figure S3.4 (cotinuation).** Predicted changes in U.S. species abundance (i.e. annual changes in the number of stems per hectare) by species-level models when setting minimum temperature and aridity index in cold and wet (blue), median climate (black), and warm and arid (red) conditions within each species' climatic niche, and setting stand development in actual, early and late stand development (see *Methods*). Points indicate mean changes in species abundance and intervals 50% uncertainty.

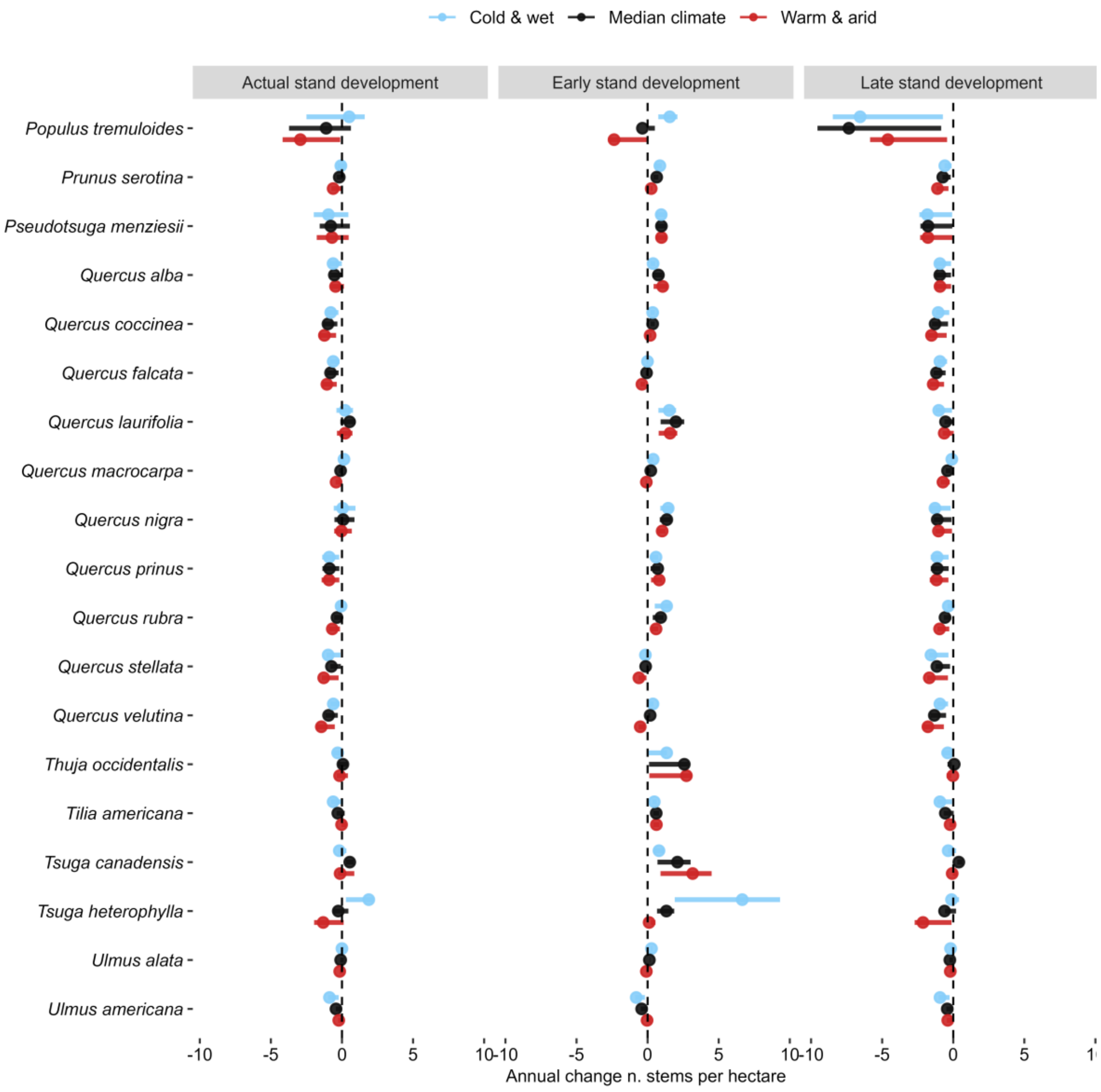


**Figure S3.4 (cotinuation).** Predicted changes in U.S. species abundance (i.e. annual changes in the number of stems per hectare) by species-level models when setting minimum temperature and aridity index in cold and wet (blue), median climate (black), and warm and arid (red) conditions within each species' climatic niche, and setting stand development in actual, early and late stand development (see *Methods*). Points indicate mean changes in species abundance and intervals 50% uncertainty.

**Table S3.1.** Percentage of species whose abundance decreases and increases in each climatic region.

| Region | Species decreasing (%) | Species increasing (%) |
|---|---|---|
| 1 (Cold & wet) | 64.4 | 35.6 |
| 2 | 69.9 | 30.1 |
| 3 (Warm & wet) | 63.0 | 37.0 |
| 4 | 56.2 | 43.8 |
| 5 | 71.2 | 28.8 |
| 6 | 71.2 | 28.8 |
| 7 (Cold & arid) | 67.1 | 32.9 |
| 8 | 71.2 | 28.8 |
| 9 (Warm & arid) | 71.2 | 28.8 |

**Table S3.2.** Mean species' climatic sensitivity. The climatic sensitivity of each species was calculated as the difference in the annual change in the number of stems per hectare between cold and wet regions, and warm and arid regions in actual, early and late stand development. Positive values indicate that in cold and wet regions species were gaining more individuals, or losing fewer individuals, than in warm and arid regions; negative values indicate the opposite. The mean climatic sensitivity of all species was calculated by averaging the mean climatic sensitivity of each species in actual, early and late stand development. Asterisks indicate statistically significant climatic sensitivities and percentages indicate the percentage of species with statistically significant climatic sensitivities in actual, early and late stand development.

| Species | Climatic sensitivity (actual) | Climatic sensitivity (early) | Climatic sensitivity (late) |
|---|---|---|---|
| *Abies balsamea* | 7.38 | 15.09 * | -0.23 * |
| *Abies concolor* | 3.93 * | 1.88 * | 4.22 * |
| *Abies grandis* | 2.2 | 6.87 * | 1.23 * |
| *Abies lasiocarpa* | 0.75 | 2.09 * | 0.93 * |
| *Acer rubrum* | 1.13 | 0.77 * | 1.06 * |
| *Acer saccharum* | -2.75 * | -3.94 * | -2.38 * |
| *Alnus glutinosa* | 10.32 | -43.16 * | 42.89 * |
| *Betula alleghaniensis* | 1.15 | 3.14 * | 0.57 * |
| *Betula lenta* | 0.97 | 5.6 * | -0.13 * |
| *Betula papyrifera* | 1.88 * | 1.65 * | 1.8 * |
| *Betula pendula* | 2.39 | -0.06 * | 3.58 * |
| *Betula pubescens* | 1.69 | 2.27 * | 0.76 * |
| *Carpinus betulus* | 0.05 | -12.68 * | 6.86 * |
| *Carya alba* | 1.02 * | 1 * | 1.03 * |
| *Carya glabra* | 0.25 * | 0.33 * | 0.22 * |
| *Carya ovata* | 0.03 | 0.5 * | -0.21 * |
| *Fagus grandifolia* | -0.07 | 2.84 * | -0.72 * |
| *Fagus sylvatica* | -3.55 | 3.88 * | -8.19 * |
| *Fraxinus americana* | 2.18 * | 2.73 * | 1.9 * |
| *Fraxinus excelsior* | -5.07 | -11.54 * | -3.31 * |
| *Fraxinus nigra* | -5.15 | -8.41 * | -3.07 * |
| *Fraxinus pennsylvanica* | -3.66 * | -6.53 * | -2.91 * |
| *Juniperus virginiana* | 0.59 | -0.07 * | 1.62 * |
| *Larix decidua* | -10.5 | -9.68 * | -16.32 * |
| *Larix laricina* | 10.32 * | 9.75 * | 10.48 * |
| *Liquidambar styraciflua* | 1.52 | 3.42 * | 0.89 * |
| *Liriodendron tulipifera* | 0.38 | -0.85 * | 1.04 * |
| *Nyssa biflora* | -0.23 | 3.24 * | 0.25 * |
| *Nyssa sylvatica* | 0.78 * | 1.15 * | 0.66 * |
| *Oxydendrum arboreum* | 1.53 * | 2.03 * | 1.42 * |
| *Picea abies* | 13.19 | -8.59 * | 25.8 * |
| *Picea engelmannii* | 0.4 * | 0.28 * | 0.39 * |
| *Picea glauca* | 2.44 | 6.76 * | -1.3 * |
| *Picea mariana* | 3.85 | 4.68 * | 2 * |
| *Picea rubens* | 2.37 | 3.42 * | 1.82 * |

| Species | Climatic sensitivity (actual) | Climatic sensitivity (early) | Climatic sensitivity (late) |
|---|---|---|---|
| *Pinus contorta* | -3.07 | -4.76 * | -2.69 * |
| *Pinus echinata* | 6.7 | 10.87 * | 6.76 * |
| *Pinus elliottii* | -4.1 | 3.63 * | -2.41 * |
| *Pinus halepensis* | 0.1 | -0.17 * | 0.19 * |
| *Pinus nigra* | 1.12 | 1.11 * | -0.05 * |
| *Pinus palustris* | 7.57 * | 7.19 * | 0.99 * |
| *Pinus pinaster* | -0.16 | -1.37 * | -0.03 * |
| *Pinus pinea* | -1.39 * | -0.53 * | -1.5 * |
| *Pinus ponderosa* | 0.18 | 0.56 * | 0 * |
| *Pinus strobus* | -2.64 | 3.28 * | -5.62 * |
| *Pinus sylvestris* | -27.86 * | -51.25 * | -2.71 * |
| *Pinus taeda* | 17.28 | 35.75 * | 10.59 * |
| *Pinus virginiana* | -2.64 | -13.39 * | 2.02 * |
| *Populus grandidentata* | 2.2 | 5.27 * | -3.42 * |
| *Populus tremula* | 12.65 | 1.75 * | 20.26 * |
| *Populus tremuloides* | 9.61 | 10.19 * | -5.07 * |
| *Prunus serotina* | 1.5 * | 1.68 * | 1.42 * |
| *Pseudotsuga menziesii* | -0.39 | -0.03 * | -0.06 * |
| *Quercus alba* | -0.48 | -1.89 * | -0.03 * |
| *Quercus coccinea* | 1.27 * | 0.51 * | 1.35 * |
| *Quercus faginea* | 1.05 * | 0.68 * | 0.5 * |
| *Quercus falcata* | 1.23 * | 1.08 * | 1.3 * |
| *Quercus ilex* | 1.43 * | 0.91 * | 1.81 * |
| *Quercus laurifolia* | -0.03 | -0.17 * | -1.08 * |
| *Quercus macrocarpa* | 1.61 * | 1.4 * | 1.81 * |
| *Quercus nigra* | 0.22 | 1.15 * | -0.66 * |
| *Quercus prinus* | 0.04 | -0.63 * | 0.13 * |
| *Quercus pyrenaica* | 0.4 | 1.47 * | -1.61 * |
| *Quercus robur* | 23.83 | 16.99 * | 25.77 * |
| *Quercus rubra* | 1.73 * | 2.08 * | 1.72 * |
| *Quercus stellata* | 0.87 | 1.28 * | 0.33 * |
| *Quercus velutina* | 2.37 * | 2.51 * | 2.37 * |
| *Thuja occidentalis* | -0.42 | -3.97 * | -1.05 * |
| *Tilia americana* | -1.68 | -0.42 * | -2.04 * |
| *Tsuga canadensis* | -0.17 | -6.71 * | -0.8 * |
| *Tsuga heterophylla* | 5.05 * | 10.31 * | 3.16 * |
| *Ulmus alata* | 0.43 | 1 * | 0.05 * |
| *Ulmus americana* | -1.82 * | -2.12 * | -1.51 * |
| Mean | 1.33 | 0.21 | 1.71 |
| Percentage | 32.9 | 100 | 100 |
